# The impact of second language immersion: Evidence from a bi-directional longitudinal cross-linguistic study

**DOI:** 10.1101/2020.09.25.311944

**Authors:** Henry Brice, Stephen Frost, Atira S. Bick, Peter J. Molfese, Jay G. Rueckl, Kenneth R. Pugh, Ram Frost

## Abstract

Brice et al. (2019) presented data from the first epoch of a longitudinal study of the neurobiological underpinnings of first-language (L1) and second-language (L2) processing. Results showed a similar network of activation for reading across L1 and L2, as well as significant convergence of print and speech processing across a network of left-hemisphere regions in both L1 and L2 with greater activation and convergence for L2 in anterior regions, and greater activation and convergence for L1 in posterior regions of the reading network. Here, we present the first look at longitudinal changes in these effects. L2 showed relatively few changes in activation, with some shifts in the weighting between ventral and dorsal processing. L1, however, showed more widespread differences in processing, suggesting that the neurobiological footprint of reading is dynamic, with both L1 and L2 impacting each other. Print/speech convergence showed very little longitudinal change, suggesting that it is a stable marker of the differences in L1 and L2 processing.

## Introduction

The neurobiological underpinnings of literacy skills have been the object of much study over the last three decades, with a growing body of research converging on a general model of how visual word recognition plays out across the brain. This body of evidence, focusing mainly on first-language (L1) reading skills, has highlighted a network of interconnected left hemisphere regions, consisting of a set of temporo-parietal, occipito-temporal, frontal, and sub-cortical regions (e.g., Pugh et al., 2010, 2013; Schlaggar & McCandliss, 2007). A dorsal system, thought to be involved in mapping orthographic input to phonological and semantic properties of written words (Xu et al., 2001), encompasses the temporo-parietal parts of the network, including the angular gyrus and the supramarginal gyrus of the inferior parietal lobule (IPL), as well as the posterior aspect of the superior temporal gyrus (STG; Wernicke’s area). Developmental studies (e.g., Booth et al., 2001; Church, Coalson, Lugar, Petersen, & Schlaggar, 2008; Martin, Schurz, Kronbichler, & Richlan, 2015) have indicated that increased reading proficiency leads to increased reliance on a more lexically mediated ventral system, spanning the ventral occipito-temporal regions of the network, including the so-called Visual Word Form Area (Dehaene, Le Clec’H, Poline, Le Bihan, & Cohen, 2002) situated in the left fusiform gyrus, as well as the middle and anterior temporal gyri, and the pars triangularis of the inferior frontal gyrus (IFG).

This reading network has been shown to be remarkably consistent across different languages, irrespective of linguistic structure or orthographic system (Pugh et al., 2013, 2000; Pugh, Sandak, Frost, Moore, & Mencl, 2005; Rueckl et al., 2015). Language-specific differences are primarily manifested in differential weighting of this common network, with some subcomponents more heavily involved in one language than another, depending on the specific linguistic and orthographic characteristics of the language in question, e.g. orthographies with more opaque mappings between written and spoken forms tend to show greater reliance on the ventral system (Bolger, Perfetti, & Schneider, 2005; Paulesu et al., 2000). This reading network is therefore taken to represent a cross-linguistic functional system of brain organisation, optimised for the task of fluent word reading (see Carreiras et al., 2014; Dehaene et al., 2015, for review and discussion).

Investigations of the neurobiological bases of second language (L2) processing have found, overall, that a very similar language network is activated for the processing of second languages (Brice et al., 2019; Das, Padakannaya, Pugh, & Singh, 2011; Perani & Abutalebi, 2005). Research has looked at how both proficiency and age of acquisition of the L2 affect reading processes (Perani et al., 1998), finding that more proficient L2s and those acquired at a younger age look more similar to the learner’s L1 (although see Das et al., 2011, for a comparison with simultaneously acquired bilingulalism). Similarity between L1/L2 pairs in terms of orthographic transparency has also been shown to affect activation (Kim et al., 2016; Liu & Cao, 2016), with more similar L1/L2 language pairs tending to have more similar activation patterns. Characteristics of L1 reading processes have been seen to carry over to an L2, both in terms of behaviour (Ben-Yehudah, Hirshorn, Simcox, Perfetti, & Fiez, 2019) and in terms of neural activation (Perani et al., 1998), despite significant differences in orthographic and morphological structure. One shortcoming of much of the current research on the neurobiology of second language literacy, however, is that it cannot fully distinguish between language-specific differences in processing, and between differences inherent to first and second languages.

In a recent study (Brice et al., 2019), we aimed to disentangle these two factors by investigating parallel cohorts of native Hebrew speakers immersing in English in the U.S., and native English speakers immersing in Hebrew in Israel. Our aim was to longitudinally track behavioural and neural changes across two years of immersion and examine to what extent the neural footprint of L2 literacy differs from that of an L1 while comparing two languages that are highly contrastive in both their linguistic and orthographic characteristics. The longitudinal design allowed us to specifically track how changes in processing and activation in both languages evolve over the course of L2 experience.

Although both English and Hebrew have relatively opaque alphabetic writing systems (see R. Frost, Katz, & Bentin, 1987), the morpho-syntactic structures of the language and their respective orthographic representations differ considerably; they utilise different alphabetic scripts in order to encode languages with fundamentally different morpho-phonological structures. Hebrew has a Semitic root-and-template morphological system, with verbs and nouns being comprised of two interleaved morphemes. This non-concatenative morphology leads to a mental lexicon that is structured differently from those of speakers of more common concatenative systems, which linearly append affixes to base morphemes (Deutsch, Frost, & Forster, 1998; R. Frost, Forster, & Deutsch, 1997; Velan & Frost, 2011). Coming from different language families, English and Hebrew also have few cognates, which have been shown to influence second language learning and vocabulary acquisition (X. Chen, Ramirez, Luo, Geva, & Ku, 2012, Tytus & Rundblad, 2017). The statistical orthographic properties of written Hebrew have been shown to affect a range of reading phenomena such as form priming and letter transposition effects (Lerner, Armstrong, & Frost, 2014; Velan, Deutsch, & Frost, 2013; Velan & Frost, 2011). The morpho-orthographic properties of the two languages have also been shown to affect neural processing, with somewhat more dorsal activation for English and more ventral activation for Hebrew, as well as an interaction with different levels of linguistic structure: semantic and morphological processing modulate each other in English but are more independent in Hebrew (A. Bick, Goelman, & Frost, 2008; A. S. Bick, Frost, & Goelman, 2010; A. S. Bick, Goelman, & Frost, 2011).

The choice of Hebrew and English as contrasting languages allows us, therefore, to examine second language acquisition of two languages that, despite both having opaque alphabetic orthographies, differ in alphabet, morphological structure and in the reading strategies that they optimally employ. The symmetric design in which each language is both L1 and L2 allows us to tease apart language-specific effects of English vs. Hebrew from more general effects of L1 vs. L2. This enables us to differentiate between changes associated with developing efficient and effective reading strategies specific to a given language’s morphological and orthographic characteristics, and changes associated with gaining proficiency in a second linguistic system when the initial L1 fluency has already been achieved.

In Brice et al., (2019) we focused on evidence from the first time-point (Epoch 1; Ep1) of L2 immersion. We showed that, overall, L2 processing resulted in greater activation in anterior regions of the language network, while L1 showed greater activation in the dorsal pathway, specifically the inferior parietal cortex and superior temporal regions. This effect was seen for both cohorts, affirming that it is an L1/L2 effect and not a result of the differences between Hebrew and English. This led us to hypothesize that if the neurobiological underpinnings of processing L2 shift with time to resemble L1, then the final epoch of the immersion period (Ep2) should show a decrease in activation in the anterior regions and perhaps an increase in activation in the posterior dorsal pathways.

Another important metric that Brice et al. (2019) reported on was the convergence of the neural networks of print and speech processing. Research by Preston and colleagues (2016) has probed this convergence in the context of L1 literacy acquisition, based on the understanding that efficient visual word processing must utilise pre-existing linguistic knowledge, leading to a functional convergence across modalities in order to achieve reading proficiency (Braze et al., 2011). Preston et al. (2016) found that the extent of overlap in the activation associated with the processing of print and speech in L1 in seven-year olds was a strong predictor of the development of reading abilities two years later. These findings suggest that successful reading acquisition for L1 results in the reorganization of the oral language networks in the brain into amodal reading related systems (Chyl et al., 2018; S. J. Frost et al., 2009; Marks et al., 2019; Shankweiler et al., 2008). In a recent cross-linguistic study (Rueckl et al., 2015), the extent of overlap between print and speech processing in the left hemisphere in adults was shown to be a hallmark of fluent reading across four languages, with a remarkably similar footprint of convergence spanning very different language and writing systems (English, Hebrew, Chinese, and Spanish).

Our first examination of print/speech convergence in L2 reading at Ep1 (Brice et al., 2019), demonstrated that the overlap between print and speech processing indeed covered regions of interest similar to those reported by Rueckl et al. (2015), but differed somewhat in the extent of convergence between L1 and L2: Greater overlap was seen for L1 in posterior parietal cortex, associated with more automatic language processing, but more overlap was seen for L2 in inferior frontal regions associated with more effortful processing. This led us to hypothesize that print/speech overlap may demonstrate longitudinal change, with L2 convergence looking more like L1 convergence as L2 proficiency increased.

With the longitudinal data collection now finished, the present paper addresses these predictions, by examining the group-level longitudinal changes in the measures reported on from Ep1. Our longitudinal tracking provides us the unique opportunity to focus on a number of important theoretical questions: **First**, Do the neurological underpinnings of literacy in L2 become more similar to those in L1 with increasing proficiency, or are such differences as were seen at Ep1 longitudinally stable? **Second**, how does longitudinal immersion in an L2 impact the neurobiological underpinnings of processing L1? Are there differences in processing between L1 before immersion (at Ep1) and following two years of immersion in L2 (Ep2)? **Finally**, can differences in the processing of each language be tied to specific characteristics of the orthographic and linguistic aspects of that language, or to the order in which they are acquired? We approach these three questions first in terms of the neural activation associated with processing L2 and then by examining the convergence of print and speech processing.

## Method

Native Hebrew speakers living in the U.S. and native U.S. English speakers living in Israel, received a functional magnetic resonance imaging (fMRI) scan, alongside a battery of behavioural measures of language processing, upon entry (Ep1), and again after two years of immersive experience (Ep2). During the brain scan they performed a semantic judgement task on printed and auditory words and pseudowords, in both L1 and L2. The aim of the semantic judgement task was to ensure full processing of the target words, in order to obtain maximal activation of all levels of linguistic analysis during word reading. In order to ascertain that L2 processing would involve full linguistic analysis, and that word recognition would be achievable at Ep1, participants were selected to have moderate proficiency in their L2, as explained below. The word stimuli chosen for the in-scanner task were selected to fit this level of L2 proficiency. In addition, we utilised a battery of behavioural measures of both language skills and general cognitive abilities to track the process of L2 acquisition, the individual level of L1 proficiency, and to ensure that the two cohorts were more or less equivalent in terms of general cognitive abilities.

In order to achieve sufficient power, we aimed to have 40 subjects providing full data in each cohort, this based on doubling the sample size in Rueckl et al. (2015), who examined cross-linguistic differences in the reading network utilising a very similar semantic judgement task. In addition to doubling the sample size from Rueckl, we are confident in the statistical power of this study given the longitudinal nature of the data, in which participants are able to act as their own controls.

### Participants

The English first language cohort (EL1) comprised 46 subjects, (31 women, mean age at start 23), 6 of whom did not provide Ep2 data. The Hebrew first language (HL1) cohort comprised 56 subjects (20 women, mean age at start 22), of whom 16 did not provide Ep2 data. Thus, 40 subjects in each cohort participated in all stages of the study, for a total of 80 subjects. The EL1 cohort all were recent immigrants to Israel from the U.S., recruited in Jerusalem, Israel, and all were participating in a Hebrew language course and/or working or studying in Hebrew. The HL1 cohort were recent Israeli immigrants to the U.S., recruited in New York, and were all working and/or studying in English. All participants reported either normal or corrected-to-normal vision, and had no diagnosis of dyslexia, dysgraphia, ADD or ADHD. Informed written consent was obtained from all participants before participation. The study was approved by the IRBs of Yale University and The Hebrew University of Jerusalem.

An important note: Certain differences in cohorts were naturally expected in extent of initial proficiency in L2. Whereas English is taught in schools in Israel and English language content is common, the reverse is not necessarily true (albeit most English L1 participants had some exposure to Hebrew script given some Jewish background). None of the HL1 cohort, however, reported use of English at home, and all of them self-reported having Hebrew as a single mother tongue. Thus, although both cohorts had some prior L2 experience, primarily with their respective L2 scripts, neither cohort had had significant L2 immersion at Ep1. For the overarching goal of the study, this was not an overly major concern, as each participant serves as their own control, providing measures of their improvement in L2 over time relative to their own baseline.

### Methods

Subjects participated in a battery of tasks measuring L1 and L2 abilities at both epochs, as well as a number of general cognitive abilities. The tasks were chosen to cover a broad spectrum of language processing, production, fluency and comprehension measures in order to provide a comprehensive picture of participant’s language skills. Subjects in both cohorts participated in four language tasks in Hebrew and two in English that were chosen to cover similar and theoretically informative aspects of language proficiency in both languages. The tasks in Hebrew were:

### Printed word naming

This task monitored both speed and accuracy in word production, as measures of familiarity and fluency in reading. Sixty printed words were presented one at a time at the centre of the screen, and participants were asked to name them aloud into a microphone headset. All words were selected from word lists for beginner level Hebrew language learners at the Hebrew university. For further details see Frost, Siegelman, Narkiss, & Afek (2013).

### Pseudoword reading

This task, developed by the Israeli National Institute for Testing and Evaluation (NITE), provides normed measures for both speed and accuracy in pseudoword reading. Participants were presented with 25 pseudowords in Hebrew script, one at a time, all written with diacritic vowel marking, which provides an unambiguous mapping to phonology. Subjects were asked to read each word aloud into a microphone headset as quickly and correctly as possible. Responses were timed and marked for correctness. Scores are reported on the normed scale provided by NITE, based on Hebrew-speaking population including both typical and atypical readers (Ben-Simon, Beyth-Marom, Inbar-Weiss, & Cohen, 2008).

### Hebrew oral passage-reading and comprehension

Subjects read aloud 9 short stories of increasing difficulty each followed by five multiple-choice comprehension questions. Speed, reading accuracy, and comprehension responses were recorded. The task was stopped if the subject answered only one question correctly for two stories in a row, or until the error rate in production went above a level proscribed for each story. The measure used for this task is the total number of stories read (see Shahar-Yames & Prior, 2018; Zeltsman-Kulick, 2016 for further details).

### Cross-modal visual lexical decision with auditory priming

This task measured phonological, morphological and semantic priming as measures of awareness of linguistics structure at various levels, and of orthographic-to-phonological mapping. On each trial, an auditory word prime was played through the subjects’ headset, followed by a printed letter string representing either a word or a pseudoword. Subjects were requested to perform a lexical decision on the letter strings. There were four priming conditions—phonologically related rhyming words with one differing phoneme; semantically related words from the same semantic field (e.g. 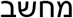, *computer* priming 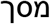, *screen);* morphologically related words derived from the same root morpheme; and a control condition with words unrelated to the targets word. The test consisted of 128 trials (64 words and 64 pseudowords, with 12 words in each priming condition). This task was included due to extensive evidence that priming measures are a reliable marker of Hebrew reading skills, tapping into morpho-orthographic decomposition of target words. See Frost et al. (2000), for details.

The tasks in English included:

### Test Of Word Reading Efficiency 2^nd^ edition

(TOWRE-2; Torgesen, Wagner, & Rashotte, 2012). This task provided measures of word and pseudoword reading. Participants were presented with a list of 104 words of decreasing frequency and increasing length, and asked to accurately read aloud as many of them as they could in 45 seconds. Following this, subjects were presented with 63 pseudowords of increasing length, and were asked to read aloud as many as they could in 45 seconds. The measure for the TOWRE is the percentage of words and pseudowords read correctly within the time limit.

### Gray Oral Reading Test 4^th^ edition

(GORT-4; Wiederholt & Bryant, 2001). This test measures fluency (a composite score of speed and accuracy) in oral paragraph reading, as well as passage comprehension. For the GORT-4, subjects were tested with 14 stories. Similar to the Hebrew oral passage reading described above, subjects read short paragraphs aloud, and then answered comprehension questions. The test provides normed scores for both fluency (speed and accuracy of reading) and comprehension.

All subjects also participated in the **Multilingual Naming Test** in both English and Hebrew (MINT; Gollan, Weissberger, Runnqvist, Montoya, & Cera, 2012) measuring accuracy in naming of a set of culturally-normed picture stimuli. Subjects were presented with 68 pictures, one at a time, on the computer screen. The frequency of the names decreases from highly frequent (e.g. *dog* and *hand*) to highly infrequent (e.g. *porthole* and *anvil*). Pictures remained on the screen until the subjects responded with a name, or said that they did not know the correct name. If an incorrect name was used, they were asked if they knew any other name before continuing to the next picture. Subjects were marked for accuracy in naming.

Finally, the in-scanner task, described below, provided measures of accuracy and speed at both epochs.

In addition to these measures of linguistic proficiency, subjects were given a number of general cognitive measures, to ensure that the cohorts did not significantly differ in their overall general abilities. These included a forward and backward **digit span task** measuring verbal working memory; **Raven’s Advanced Progressive Matrices** (Raven & Court, 1998) as a test of fluid intelligence (number of correct answers out of 36); and a **task switching** paradigm testing executive functions, in which they switched tasks every other trial, between deciding if a visually presented digit was odd or even, and whether a visually presented letter was a consonant or a vowel.

## Behavioural results

All L2 language measures (other than pseudoword reading in the HL1 cohort) showed numeric improvement between Epochs 1 and 2 (see supplementary material for tables of all tasks), and all tasks showed statistically significant improvement in at least one measure, except for the in-scanner task for the HL1 cohort, which showed a ceiling effect. Importantly, most of the Ep2 behavioural outcomes were within 1 standard deviation of the results obtained from the cohort for whom the tasks were L1, showing that L2 skills were approaching the normal range for L1 proficiency by the end of the study. We focus in here on the group-level effects, as a full investigation of the many potential individual differences is beyond the scope of this paper. Individual differences will be fully investigated in further research.

Table I presents results for the in-scanner task by cohort, L1/L2 and epoch. As can be seen, overall both cohorts had high accuracy in both L1 and L2 at both time points, confirming that the task difficulty was gauged appropriately for our participants. Both measures were analysed with an analysis of variance with the effect of cohort, time, L1/L2, and all interactions. A significant effect was seen in RT for L1/L2 (F(1,290)=48.0, p < .001), with L2 showing slower responses than L1. A marginal effect was seen for time (F(1,290)=3.65, p = .057), with faster response times at Ep2. An interaction was seen between cohort and L1/L2, (F(1,290)=11.44, p < .001), with EL1 showing slower L2 responses than the HL1 cohort. This was not unexpected, given that, as discussed, higher L2 proficiency was expected in the HL1 cohort, who had more and earlier exposure to the L2 than the EL1 cohort. The analysis of accuracy showed only an effect for cohort, with the HL1 cohort having higher overall accuracy (F(1,290)=13.8, p < .001). The lack of effects in accuracy was not unexpected, given the design of the in-scanner task, discussed below, which lead to very high overall accuracy, and something of a ceiling effect.

**Table I.** Results in the in-scanner task, in percentage of correct responses and mean RT for correct responses, by cohort, L1/L2 and epoch. Standard deviations in parentheses.

| Cohort | Epoch | Accuracy (% correct) |  | RT (ms) |  |
| --- | --- | --- | --- | --- | --- |
|  |  | L1 (Acc) | L2 (Acc) | L1 (RT) | L2 (RT) |
| English L1 | Ep1 | 93.3 (10.8) | 91.3 (12.4) | 1132 (198) | 1387 (254) |
|  | Ep2 | 93.7 (10.8) | 92.0 (11.1) | 1108 (177) | 1348 (256) |
| Hebrew L1 | Ep1 | 96.7 (2.3) | 95.7 (3.7) | 1143 (173) | 1245 (205) |
|  | Ep2 | 96.4 (4.3) | 96.1 (4.4) | 1100 (166) | 1166 (208) |

Table II presents the general cognitive measures for each cohort. No significant differences were found between the groups on these general measures.

**Table II.**
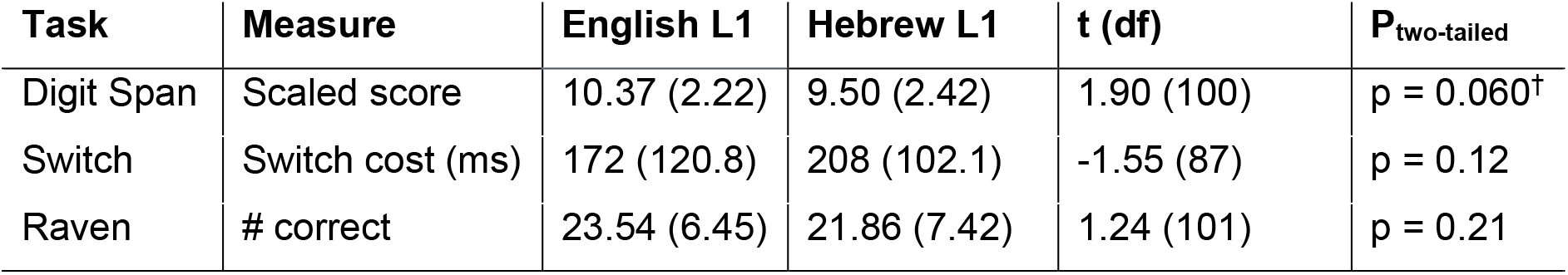
General behavioural measures for both cohorts. Standard deviations in parentheses. Degrees of freedom differ per task as not all participants had complete datasets. Digit span is translated to age-appropriate scaled scores, with a population mean of 10 and SD of 3. Raven is the number of correct responses out of 36. The switch cost is the difference in RT to trials in which the task remained the same and trials where the task switched.

These behavioural results show that participants in both cohorts improved in their L2 proficiency and fluency across the two years of the study, and approached (the lower end of) the normal range for L1 speakers. Cohorts were comparable on general cognitive skills, including verbal working memory, GI, and executive functions.

## Neuroimaging task

### Methods

To elicit activation in response to stimuli in both visual and auditory modalities and ensure full lexical access in both languages, we used a semantic categorisation task, with 3 crossed variables: L1/L2, modality (print, speech), and stimulus type (living, non-living, pseudoword) for a total of 12 stimulus conditions. Stimuli were chosen to be frequent and familiar for early L2 learners in both languages, to ensure that accuracy would be high, and that lexical access would be achieved for word targets. Subjects were asked to make semantic judgments (living/non-living) on printed and spoken isolated single words and pronounceable pseudowords in both English and Hebrew, and responded with a yes/no button press. Subjects were instructed to respond “no” (i.e., non-living) to pseudowords. Responses were recorded using an MRI-safe fiber-optic response box held in the participants’ right hand. In each of 10 functional imaging runs, event types were segregated into 4 one-minute long blocks: 1) spoken English stimuli, 2) printed English stimuli, 3) spoken Hebrew stimuli, and 4) printed Hebrew stimuli. Each block contained both real words and pseudowords. Block order was pseudorandomized across runs; in each run, the first two blocks were either both in English or both in Hebrew; the last two blocks were from the other language. A 16-second washout period was inserted in the middle of the run to separate the language blocks, with a fixation point shown for the final 3 seconds to alert participants that a new set of stimuli was about to begin. This was intended to encourage subjects to stay in a language-specific mode for longer periods of time, and to minimize effects of language switching. The experiment was presented using Presentation software (Version 16.0, www.neurobs.com). A screen on which the visual stimuli were presented was situated at the rear of the scanner bore, visible to the participants via an angled mirror. Across the 10 runs, 40 trials were obtained in each of the 8 non-living and pseudoword conditions; 20 trials were obtained in each of the 4 living conditions, for a total of 400 trials. See Figure 1. During functional scans, trials were presented at pseudo-random intervals, with inter-trial onset times jittered between 4 and 7 seconds. Occasional (10%) null trials were included to increase sensitivity, implemented as longer ITIs of 10-13s (Friston et al., 1999).

**Fig. 1.**
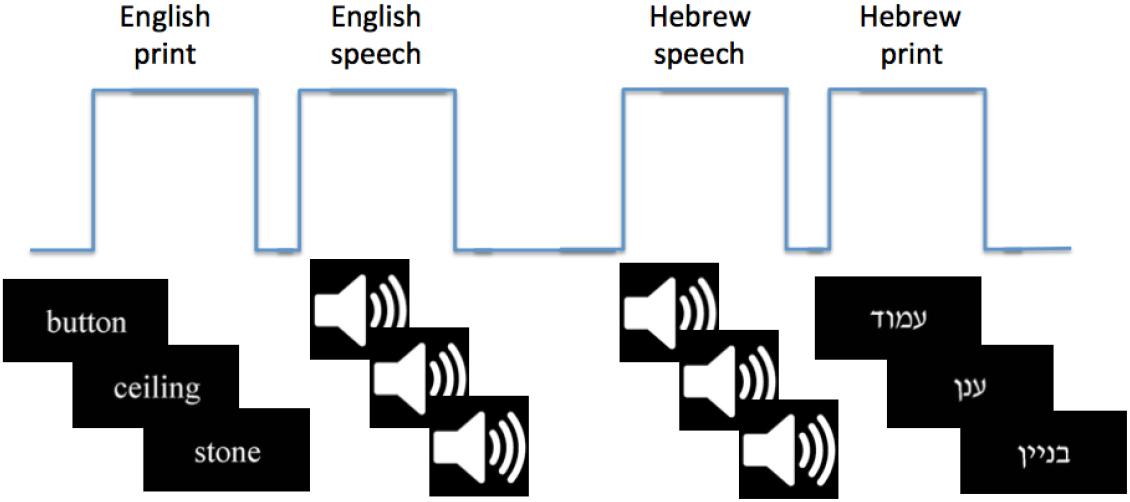
In scanner task design

**Fig. 2.**
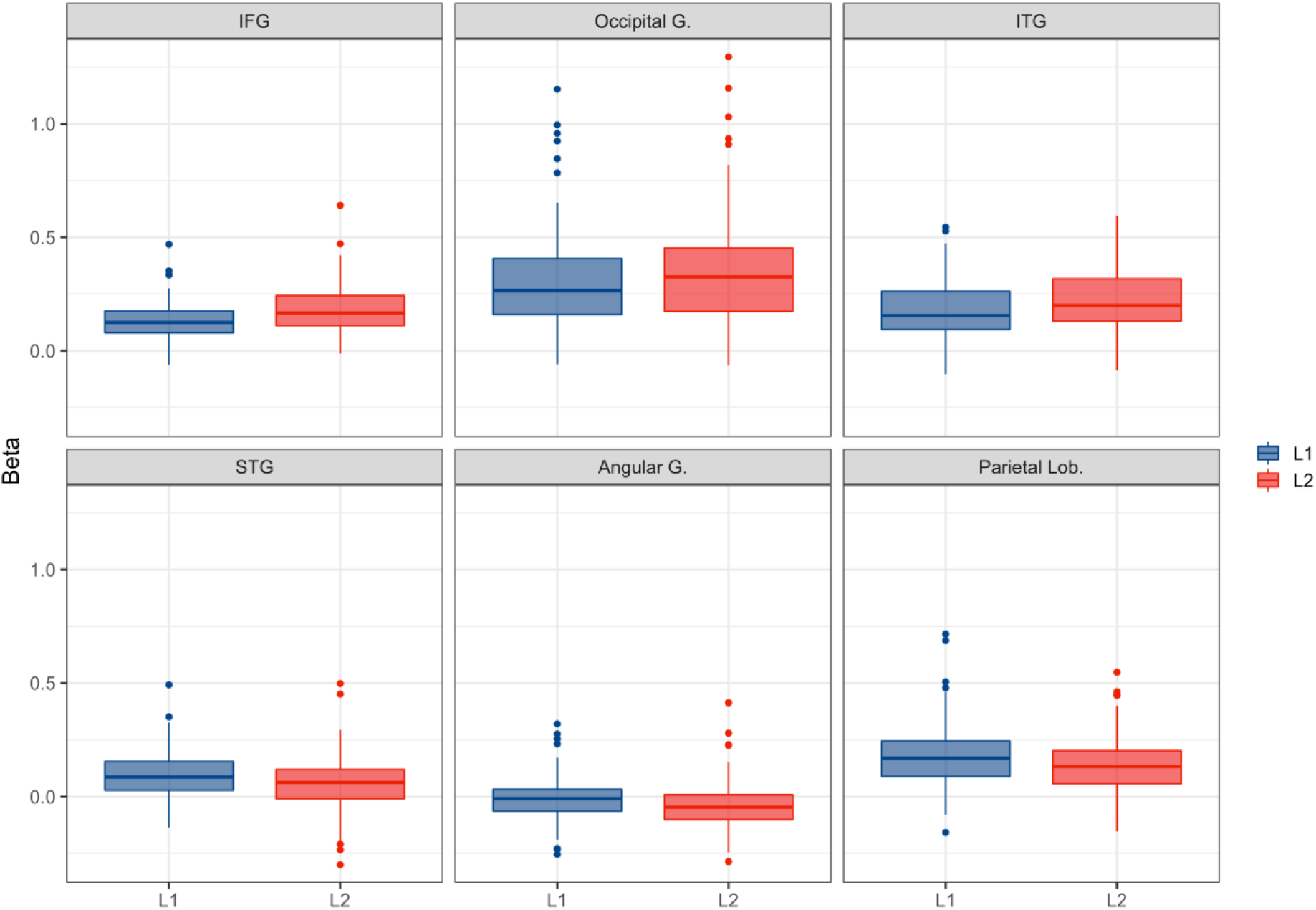
Activation for print in L1 and L2 across selected clusters in the left hemisphere. The top row shows clusters where L2 elicited greater activation, and in the bottom row L1. Similar effects were seen in the IFG, the occipital cortex, and the angular and supramarginal gyri.

Data for the HL1 cohort were collected at the New York University Center for Brain Imaging (NYU-CBI) in New York City, USA, using a Siemens Allegra 3T MRI system (Siemens AG, Erlangen, Germany) and 12-channel head coil. Data for the EL1 cohort were collected at the Hebrew University, Jerusalem, Israel, at either the Ein Karem University Hospital campus or the Givat Ram campus of the Hebrew University, using a Seimens 3T MRI system (Skyra in Givat Ram, and Trio in Ein Karem) and 12-channel head coil.

Participants were situated supine in the scanner, provided with passively noise-reducing earplugs and headphones, and stabilised with foam wedges to minimize movement. Prior to functional imaging, sagittal localizers were prescribed (matrix size = 240 × 256; voxel size = 1 × 1 × 4 mm; FoV = 240/256 mm; TR = 20 ms; TE = 6.83 ms; flip angle = 25°). Following this, an anatomical scan was acquired for each participant in an axial-oblique orientation parallel to the intercommissural line (MPRAGE; matrix size = 256 × 256; voxel size = 1 × 1 × 1 mm; FoV = 256 mm; TR = 2530 ms; TE = 3.66 ms; flip angle = 7°). Next, T2*-weighted functional images were collected in the same orientation as the anatomical volumes (32 slices; 4 mm slice thickness; no gap) using single-shot echo planar imaging (matrix size = 64 × 64; voxel size = 3.4375 × 3.4375 × 4 mm; FoV = 220 mm; TR = 2000 ms; TE = 30 ms; flip angle = 80°). Ten functional runs were acquired, each 4:24 long, for a total of 44:00 total functional running time, with 132 full-brain images each. The first six volumes within each run were discarded to allow for stabilization of the magnetic field.

### Image preprocessing and statistical analysis

Data were analysed using AFNI (Cox, 1996). Functional images were pre-processed at the single subject level by first correcting for slice acquisition time. Following this, functional images were co-registered with each subject’s anatomical images, corrected for motion using a six-parameter rigid-body transform, and then normalized to the Colin27 brain in Talairach space using a non-linear transform. These three steps were combined into a single applied transform and the data were resampled to a 3mm isotropic voxel size. Finally, all images were smoothed to the same final 8 mm smoothness level (see below).

At the subject level, data were submitted to a multiple regression analysis with nuisance regressors representing a) run-to-run mean differences; b) first- and second-order temporal drift within each run; and c) the six movement parameters. The standard BOLD hemodynamic response function (HRF) model was used as the regressor for trial events, resulting in 12 beta maps from each subject, one for each stimulus condition. For the purposes of the single-subject first-level analyses, conditions were coded as English/Hebrew, to ensure that the time-course analysis was functionally similar across subjects. Given the primary importance of L1/L2 contrasts, for the purposes of the group-level analysis this was recoded in AFNI as L1/L2 by subject.

Subject-level beta maps were used in a group-level multivariate model to examine the group-level effects of subject cohort, stimulus conditions, and their interactions (3dMVM, Chen et al., 2014).

To control for cross-scanner differences, we implemented the procedures recommended by the fBIRN consortium for multi-site scanning (Glover et al., 2012) as follows: Data from each subject were smoothed to the same final smoothness level of 8mm using an iterative procedure (3dBlurToFWHM). Then, images were scaled to percent-signal change values.

Due to the complexity of the design, and the theoretical importance of some of the higher-order interactions, groupwise statistical maps were thresholded at a relatively liberal voxelwise threshold of p = 0.005 for the purposes of the main activation effects. To control for family-wise error rates, Monte Carlo simulations were performed (3dClustSim; 10,000 iterations) using all brain voxels within the TT_N27 template brain mask, and using the spherical autocorrelation function (ACF; Cox, Chen, Glen, Reynolds, & Taylor, 2017) parameters concerning the error time series (performed in response to the recommendations regarding cluster correction in fMRI research; Eklund, Nichols, & Knutsson, 2016). The minimum cluster threshold at a p = 0.005 threshold, for a corrected alpha level of p = 0.05, was 55 voxels (3mm isotropic). Given our a-priori interest in the regions comprising the reading network, we also report where relevant on a number of clusters of activation in these regions that failed to reach the cluster threshold for significance.

## Results

Our experimental design allows for a very large number of potential contrasts. Rather than launching an exploration of all possible contrasts, and providing post-hoc justifications for those that turned out significant, we opted for a theoretically-driven approach and focused on contrasts reflecting the main theoretical questions that were at the basis of the present research project. Thus, we first looked at the primary differences in print and speech processing in L1 and L2 across both timepoints—this in light of our first theoretical question regarding overall differences between L1 and L2 processing. We then focused on the question of longitudinal changes in both L1 and L2, starting with overall activation for both print and speech. We further explored lexicality effects as a signature of language proficiency, and their evolution following immersion in L2. Finally, we focused on the extent of the overlap in neural activation across a network of ROIs canonically termed the reading network, which has been shown to reflect a signature of acquired literacy in L1 (Preston et al., 2016; Rueckl et al., 2015).

### L1 and L2 processing of print and speech

We looked first at overall differences in activation for L1 and L2, in each modality separately. Given that both English and Hebrew have relatively opaque orthographies, reliance on ventral processing is to be expected in proficient readers (e.g. Paulesu et al., 2000), and thus of particular interest is whether L1 and L2 differ in their utilisation of the dorsal and ventral pathways for the processing of print. As discussed, given the parallel cohorts of English and Hebrew speakers in our study, we can be confident that L1/L2 differences are indeed a consequence of differences in first and second language processing, whereas differences between Hebrew and English will be seen as interactions with cohort. Given our theoretical interest in print/speech integration, we also investigated differences in speech processing.

Activation associated with print stimuli showed a number of L1/L2 related differences. Greater activation was elicited for L2 (L2>L1) across much of the brain, particularly in the ventral system, including the left IFG (pars triangularis and opercularis), and spreading into the pre-central gyrus; the left posterior inferior temporal gyrus; and the right inferior and middle occipital gyri (this effect was also seen in the middle occipital gyrus in the left hemisphere, but failed to reach the cluster threshold for significance). See Table III for a list of all language differences in print.

**Table III.**
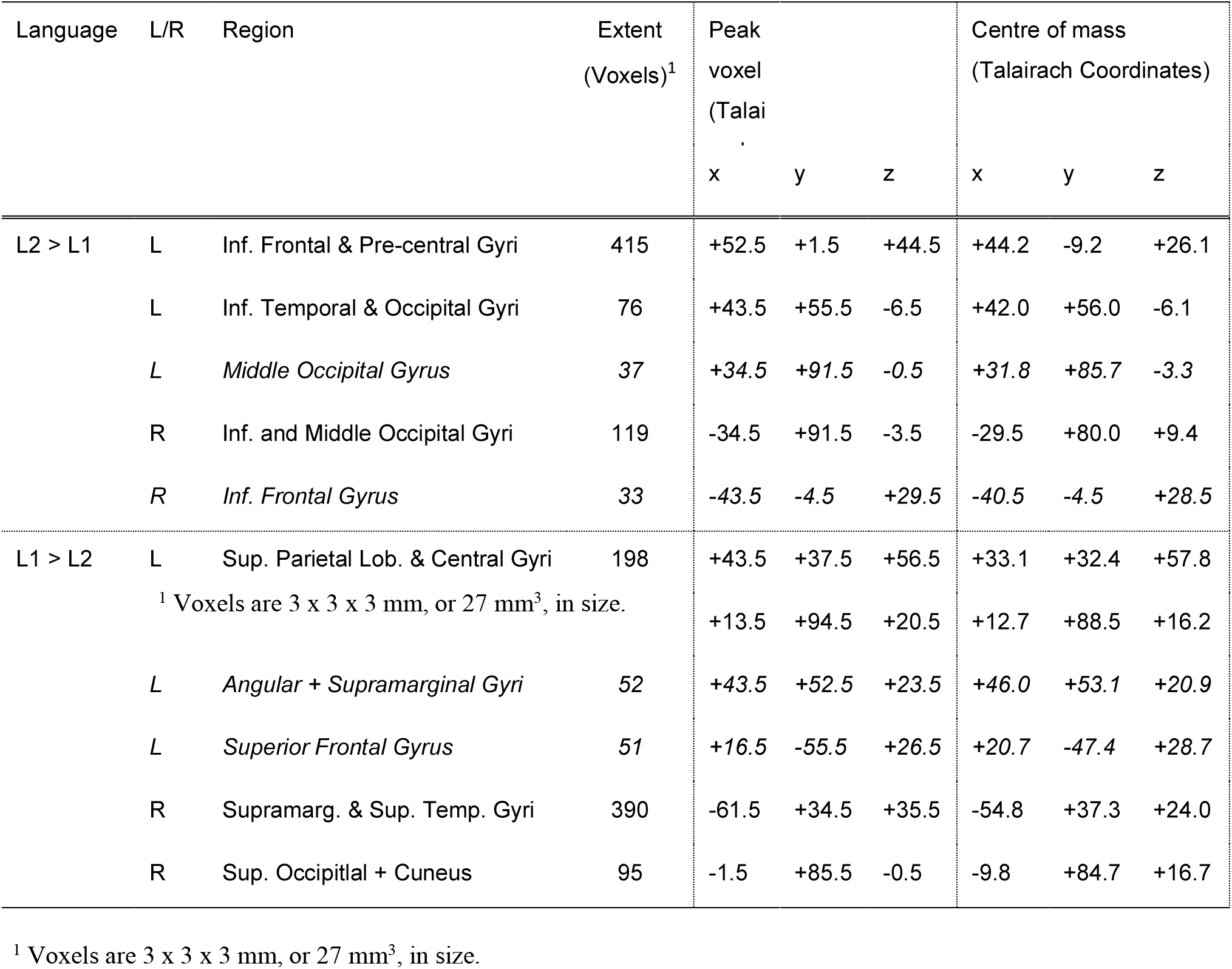
L2-L1 differences in print. Rows in italics failed to reach the cluster threshold for significance.

L2 elicited less activation than L1 (L1 >L2) in a number of dorsal regions. This effect was seen in the left superior parietal cortex and the bilateral superior occipital gyrus, as well as in the right angular and supramarginal gyri, this latter effect also seen in the left hemisphere, but failing to reach the cluster threshold.

In speech fewer differences were seen for L1/L2, with L2 showing greater activation than L1 in the bilateral IFG, and in the left hemisphere anterior STG. This shows that the frontal regions more associated with effortful processing are engaged more for the less fluent L2, as was reported in the Ep1 paper (Brice et al., 2019).

### L1 and L2 longitudinal changes in print and speech processing

Primary group contrasts of longitudinal changes in L2 and L1 word processing were conducted across the entire group for print and speech separately, in order to examine how word processing changed over the two years of exposure to the L2 for both oral and printed language, in both L1 and L2. In order not to confound the results with different semantically driven cover-task responses, for the purposes of this analysis we looked only at the inanimate word condition. Lexicality effects are explored separately below. In light of our theoretical questions, we were interested in whether differences seen at Ep1 are stable across time and whether L1 processing changes with increased exposure to L2 and reduced exposure to L1. Interactions between cohort and L1/L2 are relevant for our final question of language specificity, with opposite changes in the two cohorts able to showcase differences between Hebrew and English, rather than L1 and L2.

Activation for L1 showed significant longitudinal reduction across much of the network, with clusters of significant reduction for speech stimuli in the bilateral inferior frontal gyrus (IFG), although this failed to pass the cluster correction threshold in the left hemisphere. Effects were also seen in the anterior middle cingulate cortex, the right inferior parietal lobule (IPL) spreading into the middle occipital and angular gyri, as well as in the right cerebellum. For print, a longitudinal reduction in activation was found only in the central middle cingulate gyrus. However, it should be noted that the longitudinal trend in the IFG was modulated by cohort, an interaction that was seen also in the bilateral insula and the left STG: in all three ROIs, a reduction in activation across time was apparent for the EL1 cohort, while the HL1 cohort showed an increase in activation (see Fig. 3). In other words, L1 English showed a reduction in activation along the dorsal pathway, while L1 Hebrew showed an increase in activation in the same regions.

**Fig 3.**
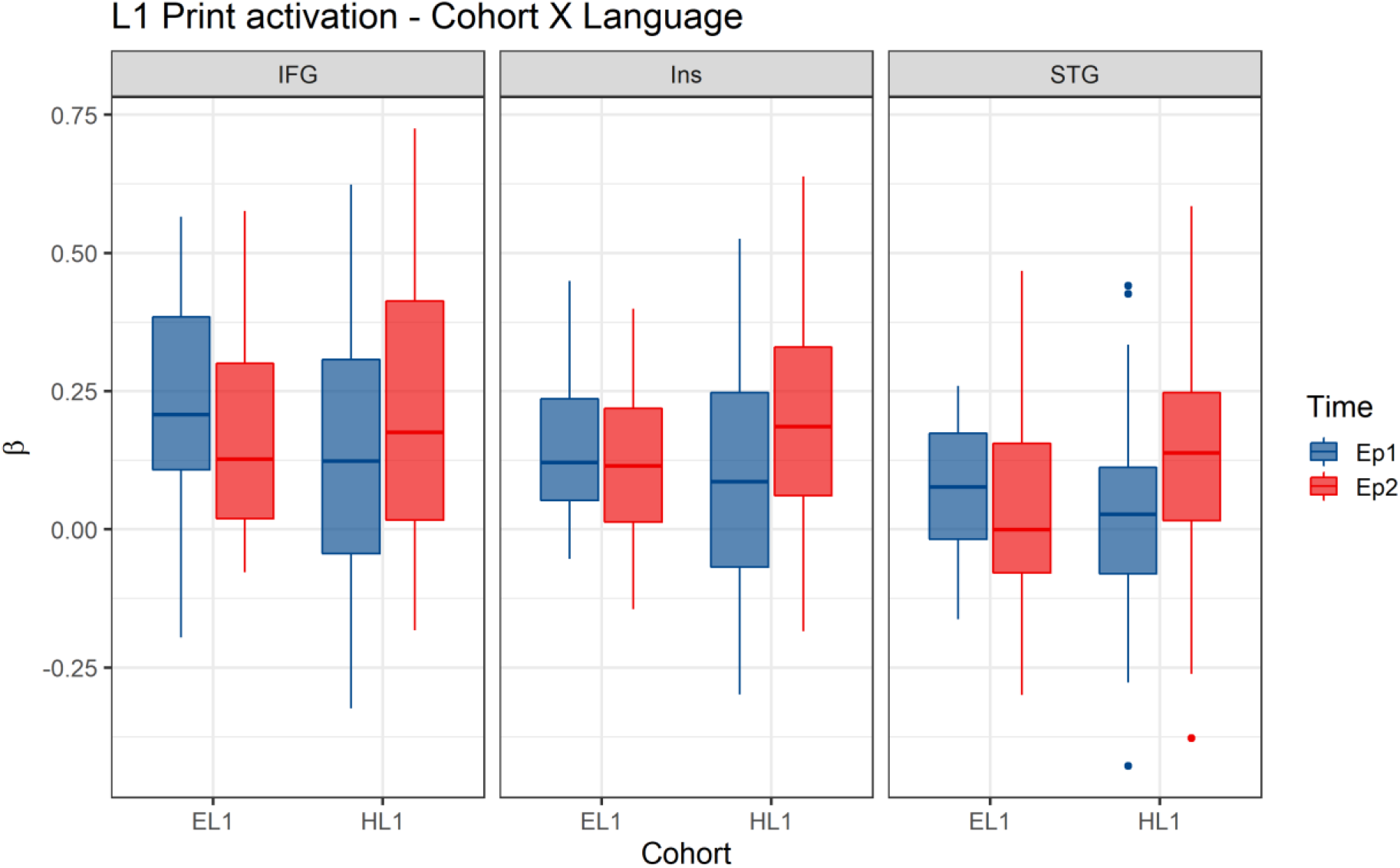
L1 print activation by cohort and epoch. The same interaction can be seen across all three clusters in the left hemisphere.

Interestingly, the processing of L2 stimuli showed less longitudinal change than that we observed for L1 stimuli. Overall, a longitudinal reduction in activation for L2 speech stimuli was found in the left middle temporal gyrus (MTG), the right post-central gyrus and the bilateral cerebellum. For print, the only effect found was a reduction in activation in the right cerebellum. Here too, however, we found a cohort by time interaction for L2 speech in both the left FFG and the right MTG, with the EL1 cohort showing an increase in activation and the HL1 cohort showing a decrease (see Fig. 4). In other words, L2 English showed a decrease in ventral activation, and L2 Hebrew showed an increase. This interaction, especially in tandem with the language effects in L1, suggest an interesting effect of learning an L2 on the processing of L1. We discuss further below.

No significant interactions were found between L1/L2 and epoch (Ep1/Ep2) in either modality. See Table IV for the list of significant clusters in longitudinal change in speech processing.

**Fig 4.**
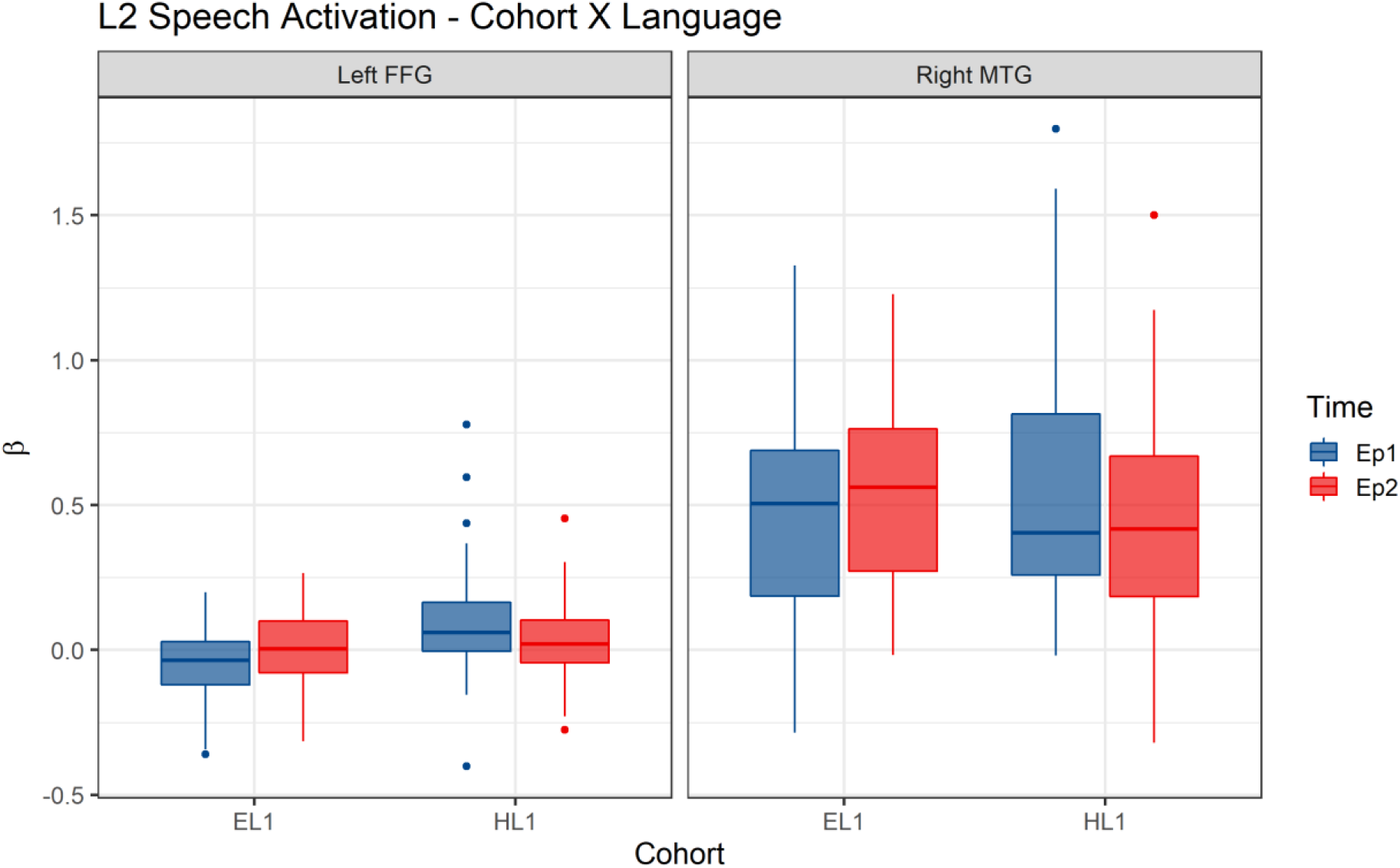
L2 speech activation longitudinal change by cohort and by language. The same interaction is seen across both clusters.

**Table IV.**
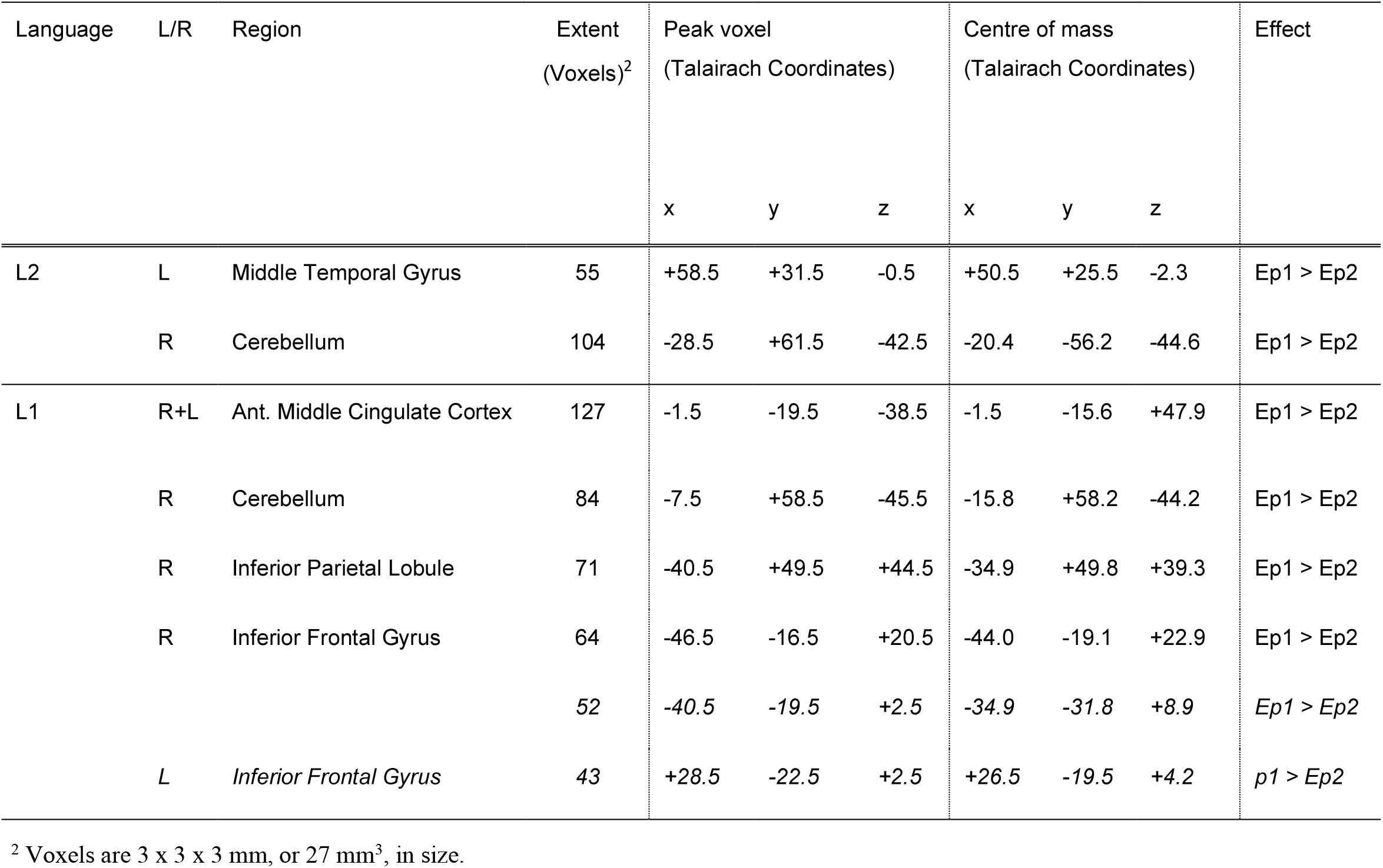
Longitudinal change in speech processing – main effects

### Longitudinal changes in lexicality effects in L1 and L2

Following the primary longitudinal contrasts, we next focused on the effect of lexicality (that is the difference in activation associated with the processing of words and pseudowords) in each modality and in each language. Importantly, this analysis compared only the inanimate words with the pseudowords, so as to remove any confound of response type to the animacy judgement cover task. Lexicality effects such as these can show changes in the sensitivity of subjects to the lexical content of the target words. A difference in the processing of words and pseudowords shows that words are being recognised and analysed accordingly. As such, we would expect lexicality effects to be affected by the speed and ease of lexical access, which can be driven by recent exposure. Given the reduced exposure to L1 and increased exposure to L2 that our participants experienced over the course of the study as they were immersed in a new linguistic environment, we expected longitudinal change in lexicality effects.

#### Print

A three-way interaction was found between lexicality, L1/L2, and epoch in the bilateral fusiform gyrus, spreading into the inferior occipital gyri (see Fig.5). At Ep1, L1 showed an effect of greater activation for pseudowords than words. L2, however, showed a reverse effect, with word stimuli showing more activation than pseudoword stimuli. Thus, for L2, recognition of word stimuli implicated greater involvement of the ventral pathway than for pseudoword stimuli. However, by Ep2, these effects had reversed, with word stimuli showing greater activation than pseudoword stimuli in L1, but less activation in L2. This suggests that the effect of lexical content on processing in the lexically-mediated ventral stream may be being driven primarily by exposure, with the language that is getting more daily use (L1 at Ep1, but L2 at Ep2) showing greater ventral involvement for pseudowords, but the language with reduced exposure showing greater ventral involvement for words.

**Fig 5.**
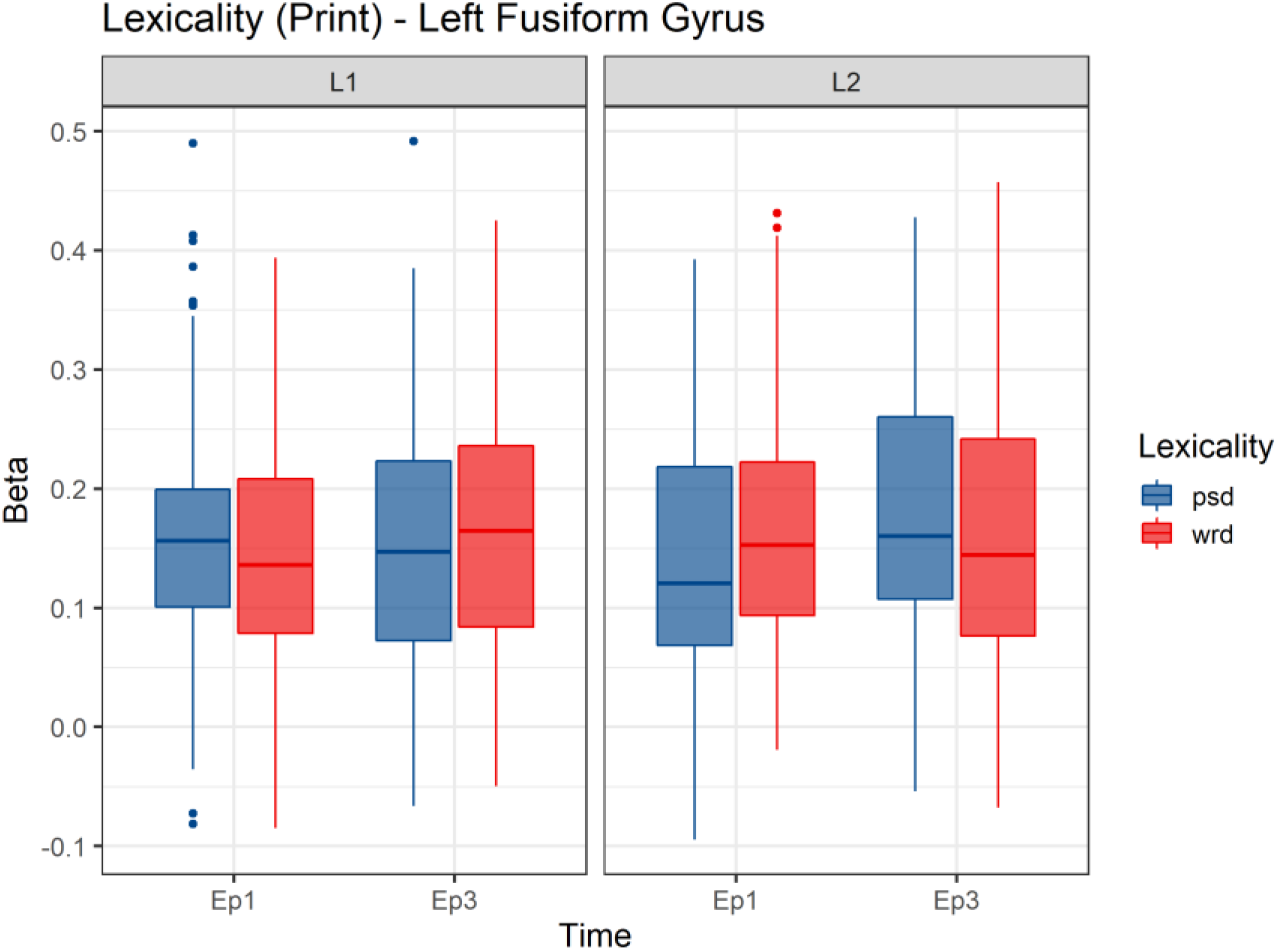
Lexicality effect in print in the left Fusiform Gyrus for each epoch in L1 and L2. Similar effects were seen in the right FFG, but smaller in magnitude.

**Fig 6.**
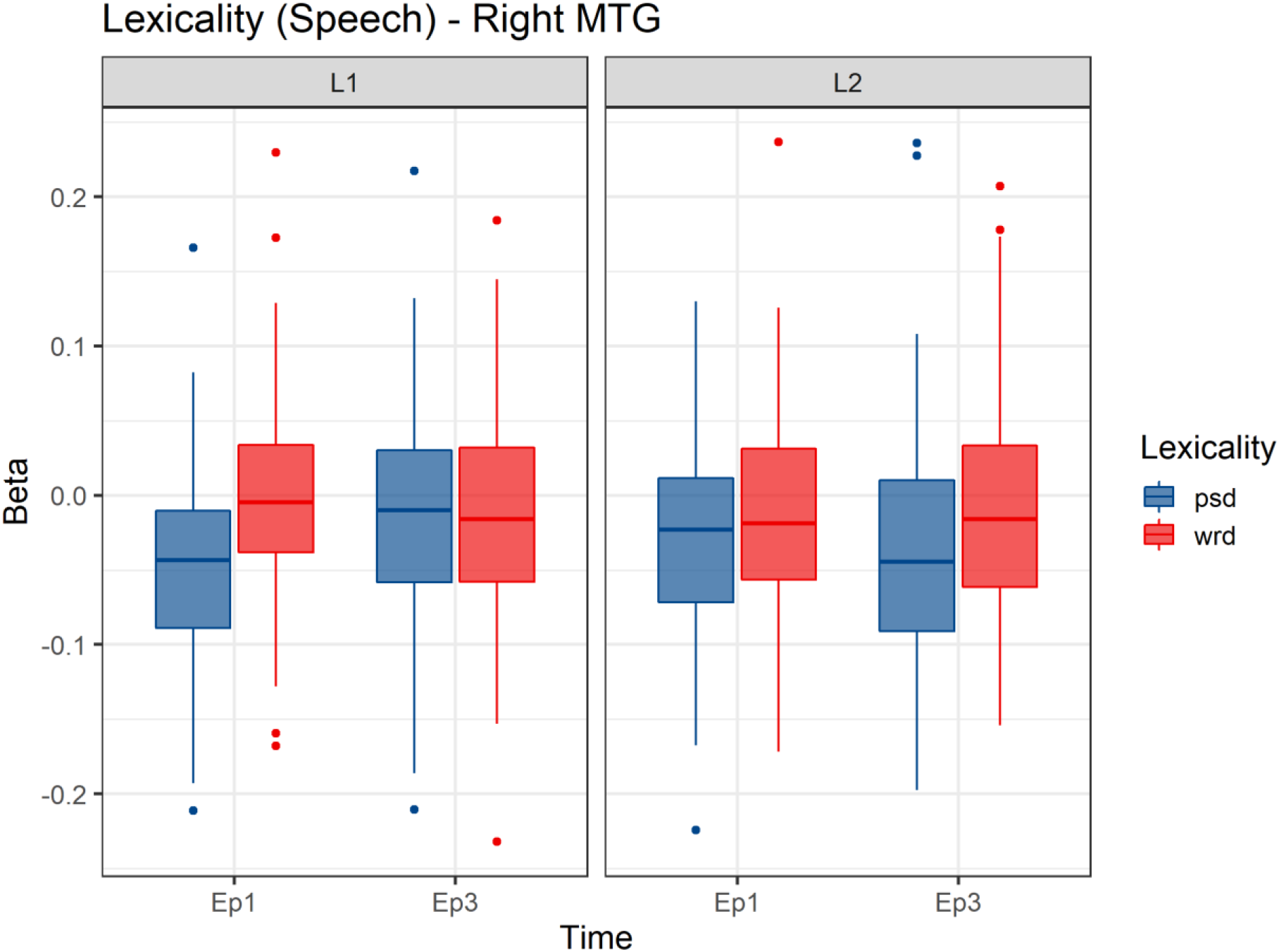
Lexicality effects in speech in the right Middle Temporal Gyrus for L1 and L2.

#### Speech

A similar three-way interaction was seen across the right posterior superior and middle temporal gyri, spreading into the angular gyrus (see Fig. 5). Here the lexicality effects were somewhat simpler, with words here showing greater dorsal activation than pseudowords at Ep1 in both L1 and L2. This effect remained stable in L2, but was greatly reduced in L1. This difference in lexicality effects as compared to those found in print could be due to modality differences or perhaps to differences in ventral and dorsal processing.

In both modalities we therefore see that early L1 lexicality effects are either reduced or reversed with the decrease in exposure, while L2 lexicality effects are either stable, or begin to show L1-like effects with time. These findings thus offer strong evidence that lexical access efficiency in both languages is affected by the two years of consistent exposure to the new L2, and reduced exposure to the L1.

### Summary – whole brain activation

The results of the whole-brain L1/L2 comparison show that the ventral system is being more strongly activated by L2 than by L1, keeping in mind that both languages are relatively opaque. The dorsal system, however, shows less activation for the L2 than the L1. This raises the intriguing possibility that, for L1 readers of an opaque orthography in L1, reliance on the dorsal system may be reduced from the start in an L2, with the more efficient ventral reading strategies developed during literacy acquisition in the L1 being carried directly over into the L2. This is an interpretation that will necessitate further investigation with additional language pairs but, if true, it suggests why L1-L2 language pairs that are similar in their orthographic opacity show more similar activation patterns (Kim et al., 2016).

Interestingly, the examination of longitudinal change shows, overall, more changes in the processing of L1 than of L2, with L1 showing a decrease in activation across a range of higher-level language processing regions, while L2 shows a decrease in activation in a smaller number of primary processing regions, primarily in speech. The relative lack of change in the activation associated with L2 processing is particularly surprising given the significant improvement seen in the behavioural measures of L2 processing in both cohorts.

One possible account for the lack of strong changes in activation associated with L2 processing is the participants’ proficiency in L2 at Ep1. As discussed, participants were initially chosen to have moderate proficiency in their L2 to allow for a reasonable behavioral assessment of L2 processing. Our findings suggest that perhaps the most dramatic changes in the neural processing of L2 occur at the very early stages of acquisition, when the most basic elements of the language are being learned, with only minor neural changes seen at later stages. Another possibility to consider, however, is that the neural changes in the processing are non-linear, with the importance of various sub-components of L2 processing plateauing or even reversing as proficiency increases. Such variable individual longitudinal differences could potentially be lost at the group level. An in-depth investigation tracking individual differences in longitudinal emersion is beyond the scope of the current paper, however, this presents an interesting future research avenue.

One of the most interesting longitudinal changes in processing is the interaction between time and cohort, which shows opposite effects in the ventral and dorsal portions of the network. Given our parallel design, interactions such as these allow us to disentangle L1/L2 effects from Hebrew/English effects. English showed a longitudinal reduction in activation in the dorsal system as L1 and a reduction in the ventral system as L2, while Hebrew showed increased dorsal activation across time as L1 and increased ventral activation as L2. We interpret this as a complex interaction between language and exposure, with increased L2 exposure affecting the processing of L1. Recalling that Bick et al. (2011) found that Hebrew shows greater ventral involvement and less dorsal activation than English, it makes sense that L2 readers will initially utilise their L1 reading strategies in reading an L2, but shift to the more optimal strategy as their L2 progresses—an increase in ventral activation for Hebrew and a decrease for English. Crucially, however, the new L2 strategies, combined with the reduced exposure to the L1, appear to be impacting the L1 in the dorsal stream, with English L1 showing less dorsal activation as the reader is immersed in Hebrew, and Hebrew L1 showing more dorsal activation due to immersion in English. This language-specific effect shows that not only do L2 reading strategies develop, but that L1 reading strategies are affected by exposure to a new L2.

The effect of lexicality, i.e. the differences between the processing of words and pseudowords, is also particularly interesting. Here we see significant changes across primary language processing regions in both modalities. The lexicality effects seen for L1 at Ep1 are reduced over time, but begin to be seen for L2 at Ep2. Thus, the driver of lexicality effects seems to be the language that is getting greater daily use, rather than overall proficiency in the language.

Overall, we see that L2 shows a very similar pattern of neural activity to L1, utilising the same networks, but weighting them according to the specific characteristics of the language in question. Interestingly, these differences seem relatively stable despite improvement in L2 fluency and exposure. The neural activity associated with L1, however, shows a broader pattern of changes, with the processing of L1 affected both by the developing processing strategies for the L2 in which the learner is immersed, and by the reduced exposure to the L1.

### Print-speech convergence

We turn next to the investigation of convergence in neural activation associated with print and speech stimuli, across a network of ROIs developed from previous research (Chyl et al., 2018; Marks et al., 2019; Preston et al., 2016; Rueckl et al., 2015). The reading network is comprised of the inferior frontal gyrus (IFG; including the pars orbitalis, triangularis and opercularis), the superior temporal gyrus (STG), the inferior parietal cortex (IPC; including the inferior parietal lobule and the angular and supramarginal gyri); and the fusiform gyrus (FFG). The four bilateral anatomical ROIs in each hemisphere were defined using neuroanatomical regions taken from the Colin27 atlas included with AFNI.

Following S. Frost et al. (2009) and Rueckl et al. (2015), we created a metric of convergence based on coactivation, this to identify the overlap between neural activity associated with processing print and speech in both L1 and L2. This was defined as the proportion of voxels within an ROI that were significantly activated (*p* < .01) for both print and speech word stimuli (independent joint probability *p* < .0001). This threshold has been previously shown to reveal reliable individual differences in coactivation in L1 that were predictive of L1 behavioural differences in both adults (S. Frost et al., 2009; Rueckl et al., 2015) and children (Preston et al., 2016). In addition, the number of voxels across the whole brain activated at *p* < 0.01 for print and for speech was computed, in order to control for individual differences in overall brain activation.

The measures of convergence were analysed using linear mixed effect models as implemented in the lme4 package for R (Bates, Mächler, Bolker, & Walker, 2015), with the lmerTest package for measuring p-values (Kuznetsova, Brockhoff, & Christensen, 2017). Models were run separately for each ROI, and included sum-coded main effects and interactions of cohort (EL1/HL1), time (Ep1/Ep2), hemisphere (R/L), and L1/L2, as well as the two whole-brain control variables for activation for print and for speech. For each model the maximal random effects model that converged was used (following the recommendations of Barr, Levy, Scheepers, & Tily, 2013) starting from by-subject random intercepts and random by-subject slopes and interactions for time, hemisphere, and language. As with the activation effects, given the complexity of the model, we report here only on the theoretically motivated effects of L1/L2, time, and their interactions with cohort. The full specification of all models and their full output can be found in the supplementary material.

### Longitudinal change in convergence

As discussed above, a theory-driven approach was taken to the examination of the convergence measure, with a focus on L1/L2 differences and longitudinal change, and their interaction with each other and with cohort. Brice et al. (2019) found that while L1 and L2 convergence was seen in similar regions of the brain, the weighting in different regions of the network differed with between the two, with greater convergence for L2 in frontal regions but greater convergence for L1 in the parietal cortex, which we interpreted as a greater reliance on more automatic processing for L1, and more effortful processing for the less proficient L2. This led us to the prediction that L2 convergence would shift with increasing proficiency, with less convergence in frontal regions, and more in the parietal cortex. We therefore examined convergence in each ROI separately (see Fig.7).

**Fig 7.**
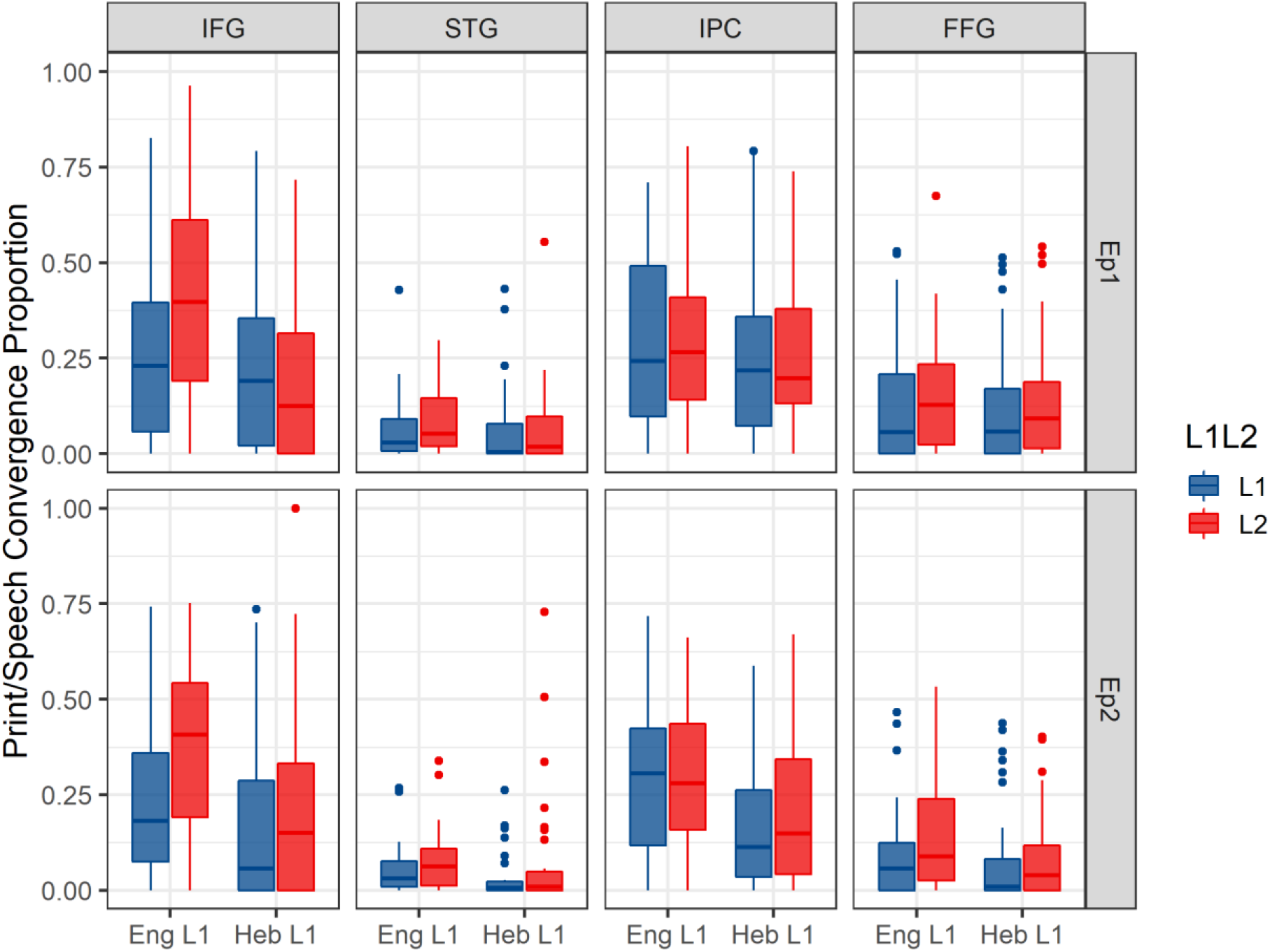
Print/speech convergence in the four ROIs, by cohort, L1/L2 and epoch.

Overlap in the IFG showed a main effect of L1/L2, with L2 showing more overlap than L1 (ß = 0.022, t(101) = 4.71, *p* < 0.001), replicating the effect found at Ep1. This was modulated by an interaction between time and language (ß = 0.008, t(2068) = 2.26, *p* = 0.024), with a surprising reduction in L1 overlap over time, while L2 overlap, which was greater in magnitude at Ep1, remained stable. This interaction was further modulated by a second order interaction with cohort (ß = 0.008, t(2068) = 2.041, *p* = 0.041), with the HL1 cohort showing greater reduction in L1 overlap. This shows that the original finding of greater overlap in L2 as compared to L1 in the IFG at Ep1, contrary to our predictions, not only persisted across time, but if anything grew even stronger with increased exposure and proficiency in L2, and reduced exposure to L1.

The STG showed a complex set of interactions. A first order interaction was found between time and L1/L2 (ß = −0.005, t(470) = −2.93, *p* = 0.004), with L1 overlap remaining stable, and with L2 showing a surprising decrease in overlap with time. However an additional interaction was seen between cohort and L1/L2 (ß = 0.010, t(98) = −3.69, *p* < 0.001), with EL1 showing more overlap for L2 than L1, and HL1 showing the reverse effect. In other words, we see a language specific effect, with Hebrew showing greater overlap than English, whether an L1 or L2, an effect which stable across time and increased proficiency in L2.

The IPC showed a main effect of L1/L2 (ß = 0.014, t(105) = −3.06, *p* = 0.003), with L1 showing more coactivation than L2, as was seen previously in the Ep1 data. This difference between the IFG and IPC shows that the balance of the overlap measure differs across the reading network, with frontal regions associated with more effortful processing showing more overlap in L2, while posterior regions associated with automatic fluent processing show more overlap for L1. Notedly, this difference also remains stable despite the increase in proficiency for L2 and the decrease in exposure to L1.

Overlap in the FFG showed only an effect for time, with a slight reduction in print-speech convergence across time (ß = −0.013, t(90) = −2.60, *p* = 0.011), however no effects or interactions were seen for L1/L2 or cohort.

### Summary – Print/speech Convergence

The analysis of the Ep1 results reported in Brice et al. (2019) showed a number of differences between languages, with greater overlap for L2 than L1 in the IFG, and greater overlap for L1 in the IPC. This was interpreted as showing a differential in overall cross-modal linguistic processing, with the more effortful processing needed for L2 causing greater overlap in frontal regions, and the more automatic processing that came with greater fluency and familiarity with L1 leading to greater overlap in more automatically associating posterior regions. These findings led to the prediction that L2 processing would show a longitudinal shift in overlap from frontal to posterior regions, with a reduction of overlap in the IFG, and an increase in the IPC, looking more like the L1 as L2 proficiency increased.

What we see in our examination of the longitudinal changes, however, is that the original findings are replicated longitudinally. If anything, the longitudinal increase in the L2>L1 effect in the IFG, alongside the cohort interaction, and the marginal interactions in the IPC, suggest a strengthening of the language differences over time. The difference in overlap between L1 and L2 is therefore a stable effect, and a potential hallmark of the difference between L1 and L2 processing.

In addition to these two ROIs in which effects were found at Ep1, an intriguing possibility is raised by the cohort by language interaction seen in the STG, with the EL1 cohort showing greater overlap in L2, and the HL1 showing greater overlap in L1. This L1/L2 difference could be due to an interaction with the orthographic characteristics of the languages, with English showing less involvement of phonological processing in reading than Hebrew, perhaps due to the greater morpho-phonological complexity of the Hebrew orthographic system (Velan & Frost, 2011). This effect is numerically more pronounced at Ep2, and although the three-way interaction with time is not significant, the effect was not evident at Ep1 (Brice et al., 2019). These two interactions occurring together however, with L1/L2 differing both by time and by cohort, are particularly interesting. Although Hebrew shows greater print/speech overlap than English in the STG, this may be modulated by recency and proficiency, as Hebrew actually shows an increase in overlap for L1 speakers across time, but a decrease in overlap for L2 speakers. This suggests that O-to-P mappings may be relied on more for processing when the language is seeing less frequent use, irrespective of their general effectiveness in the given language. Although an explicit test of this hypothesis will necessitate further study, it is nonetheless an intriguing suggestion of how differences in the orthographic system may play into the overlap of print and speech processing and the involvement of phonology in the processing of orthography and vice versa, as proficiency increases.

The results of the print/speech overlap longitudinal analysis therefore suggest that the differences in the neurological footprint between L1 and L2 are therefore mostly stable over time, despite increasing proficiency. This raises the intriguing possibility that differences in L1 and L2 overlap may be a stable neurological footprint of a second language, independent of fluency or proficiency in the L2. Although developmental evidence has shown that convergence increases as reading skills development, and also that individual differences in convergence are predictive of the development of literacy skills, it seems relatively stable in moderately proficient L2 learners. This stability of convergence in L2 is supported by recent evidence from Gurunandan and colleagues (Gurunandan, Carreiras, & Paz-Alonso, 2019), who found no differences in L2 print/speech convergence between intermediate and advanced L2 readers.

### Summary and conclusions

This study is the first large-scale study we know of to examine the neurobiological underpinnings of L2 literacy acquisition from a longitudinal perspective with parallel cohorts that allow us to disentangle L1/L2 effects from language-specific differences. We have shown that overall, the universal L1 reading network is being utilised by proficient L2 readers to read their L2. As with cross-linguistic differences in L1, the weighting of the network is affected by the specifics of the orthographic and linguistic system of the language in question, with the regularity of mappings from orthography to phonology, morphology and semantics changing optimal reading strategies

Interestingly, we have shown that the processing of the L1 is affected by a reduction in exposure to the L1, but also by the characteristics of the new L2. As new L2 processing strategies are employed, they will impact the way the L1 is processed. Indeed, L1 overall is apparently more affected than L2 by the longitudinal immersion in L2. The processing of an L1 is therefore not static, and reading strategies can shift with a change in the linguistic environment and exposure. These effects are seen primarily in the weighting of activation in the ventral and dorsal streams, with the increased morpho-phonological complexity of Hebrew leading to more overall reliance on ventral processing for those immersed in Hebrew, and immersion in English leading to more overall reliance on dorsal processing. Complementary effects were found also for lexicality, with changes to lexicality effects mostly in L1 as exposure was reduced over time. The driver of lexicality differences therefore seems to be recent daily exposure, rather than overall proficiency. This, like the primary results of the activation analysis, suggests that reading strategies are not static even within a proficient L1, but adapt to account for a new linguistic environment. Acquiring L2 proficiency is indeed impacted by pre-existing L1 skills, but pre-existing proficient L1 processing will also be affected by the acquisition of literacy in a new language. Indeed, one of the most striking findings of this study is the fact that far greater changes were seen in the processing of L1 than L2, which remained relatively stable across the two years, despite the behavioural evidence for increased proficiency and fluency in the respective L2s.

Our examination of the print-speech convergence, despite reaffirming the finding that print-speech overlap is, overall, a universal hallmark of proficient readers (Rueckl et al., 2015), found systematic and stable differences between L1 and L2 in the weighting of this overlap across the brain, with greater overlap for L2 in frontal regions, but greater overlap for L1 in posterior regions of the reading network. As proposed in Brice et al. (2019), this suggests that the measure of print-speech overlap is sensitive both to some language specific differences (as seen in the STG), but also differs between L1 and L2, with overlap being greater for L2 in parts of the ventral stream, and greater for L1 in parietal regions associated with the dorsal stream. The possibility that this is a stable hallmark of a second language is an intriguing one, suggesting that this core characteristic of reading proficiency is perhaps dependent on age at language acquisition. Print/speech convergence may be an overly monolithic measure in proficient L2 adults, despite being a predictive measure in development. Further investigations of systems-level processing or voxel-wise multivariate modelling may turn up more fine-grained differences in how L1 and L2 play out across the brain networks. We are currently investigating both these types of analysis and also measures of individual differences, the results of which will be forthcoming.

One limitation of the current study is that both Hebrew and English, despite having different alphabets and different morphological systems, both have relatively opaque orthographies. It is thus possible that the stability of the overlap differences is related to effective strategies for decoding opaque orthographies, or to the fact that subjects were moderately proficient at their L2 even at Ep1. Coming from an opaque orthography may help L2 learners utilise effective lexically mediated processing strategies in the ventral stream, without the initial reliance on dorsal processing that is a hallmark of early stages of reading development in an L1. Greater reliance on the dorsal stream is more effective a strategy for shallow orthographies, where phonological decoding is more informative for the reader (Bolger et al., 2005; Paulesu et al., 2000). This suggests that switching from an opaque to a shallow orthography, or vice versa, may show different effects from those shown here, both in the weighting of the various components of the ventral and dorsal processing streams, but also in the distribution of print-speech overlap across the brain.

Supplementary material for this paper can be retrieved from https://osf.io/cuh8n/

## Supporting information

Supplemental behavioural results

Supplemental convergence data

