## Supplemental behavioural results for "The impact of second language immersion: Evidence from a bi-directional longitudinal cross-linguistic study"

**Supplementary material – Behavioural Measures**

| Task | Measure | L1 | Epoch 1 | Epoch 2 | Δ | t(df) | $P_{two\text{-}tailed}$ |
|---|---|---|---|---|---|---|---|
| Naming | RT (ms) | 514 (70) | 839 (264) | 817 (247) | 22 | 1.307 (38) | p = 0.199 |
| | Acc. (%) | 99.3 (1.4) | 83.4 (13.4) | 88.5 (11.8) | 5.1 | 5.301 (39) | $p < 0.001^{***}$ |
| Priming | Phon. (ms) | -4.7 (62.9) | 38.1 (84.1) | 52.9 (78.2) | 14.76 | 0.704 (37) | p = 0.486 |
| | Morph. (ms) | 20.6 (38.1) | 49.1 (89.6) | 54.1 (69.6) | 5.03 | 0.025 (37) | p = 0.980 |
| | Semantic (ms) | 17.8 (42.4) | 37.4 (75.2) | 51.6 (89.7) | 14.16 | 0.543 (37) | p = 0.591 |
| | RT (ms) | 597 (109) | 839 (231) | 749 (190) | 90 | 5.700 (37) | $P < 0.001^{***}$ |
| | Acc. (%) | 96.8 (4.4) | 85.8 (12.1) | 90.0 (9.8) | 4.24 | 4.294 (37) | $p < 0.001^{***}$ |
| Oral Reading | # stories read | NA | 6.2 (2.1) | 7.3 (1.7) | 1.1 | 4.146 (33) | $p < 0.001^{***}$ |
| Pseudoword Reading | RT (z) | NA | -1.71 (2.1) | -1.40 (1.8) | 0.318 | 1.187 (33) | p = 0.245 |
| | Acc. (z) | NA | -0.27 (1.3) | -0.03 (1.2) | 0.245 | 2.275 (39) | $p = 0.028^{*}$ |
| MINT | # correct | 61.5 (2.5) | 56.4 (11.8) | 61.6 (11.1) | 5.2 | 4.546 (39) | $p < 0.001^{***}$ |
| Semantic Categorisation | Acc. (%) | 96.7 (0.2) | 91.3 (12.4) | 92.0 (11.1) | 0.68 | 0.607 (34) | p = 0.548 |
| | RT (ms) | 1145 (179) | 1393 (249) | 1353 (243) | 40 | 1.618 (34) | p = 0.115 |

**Table I** *Behavioural measures on Hebrew tests.* Standard deviations in parentheses. Note that degrees of freedom vary across tasks, due to missing values and data collection errors that were removed on a pairwise basis for each test separately. The oral reading task does not have comparable L1 measures, as different subsets of the task were used for L1 and L2 readers. Pseudoword reading is normed on native Hebrew readers (neurotypical and reading disability), with a mean of 0 and a SD of 1.

| Task | Measure | L1 | Epoch 1 | Epoch 2 | Δ | t(df) | $P_{two\text{-}tailed}$ |
|---|---|---|---|---|---|---|---|
| GORT-4 | Fluency | NA | 73.0 (24.9) | 90.8 (27.5) | 17.8 | 6.778 (35) | $p < 0.001^{***}$ |
| | Comprehension | NA | 41.2 (8.9) | 46.4 (8.1) | 5.2 | 4.665 (35) | $p < 0.001^{***}$ |
| TOWRE | Word correct (%) | 93.3 (7.4) | 74.5 (11.7) | 80.6 (8.9) | 6.1 | 4.642 (38) | $p < 0.001^{***}$ |
| | Pseudoword (%) | 86..3 (9.4) | 74.2 (13.2) | 73.3 (17.2) | -1.1 | 0.418 (38) | p = 0.678 |
| MINT | # correct | 62.7 (2.4) | 48.5 (7.0) | 54.3 (5.6) | 5.8 | 11.12 (38) | $p < 0.001^{***}$ |
| Semantic Categorisation | Accuracy (%) | 93.3 (10.7) | 95.7 (3.6) | 96.1 (4.4) | 1.4 | 0.398 (35) | p = 0.693 |
| | RT (ms) | 1121 (195) | 1235 (194) | 1164 (192) | 71 | 2.272 (35) | $p = 0.029^{*}$ |

**Table II** *Behavioural measures on English tests*. Standard deviations in parentheses. Note that degrees of freedom vary across tasks, due to missing values and data collection errors that were removed on a pairwise basis for each test separately. The GORT does not have comparable L1 measures, as different subsets of the task were used for L1 and L2 readers.
