## Supplemental convergence data for "The impact of second language immersion: Evidence from a bi-directional longitudinal cross-linguistic study": Print_Speech_Overlap.R

```
require(lme4)
require(lmerTest)

# IFG analysis ---------------------
IFG_ps_all <- readRDS("IFG.Rda")
IFGall_lme <- lmer(data=IFG_ps_all, Overlap ~ cohort*time*Hemisphere*L1L2+print_base+speech_base + (1+L1L2|subject))
summary(IFGall_lme)

# Linear mixed model fit by REML. t-tests use Satterthwaite's method ['lmerModLmerTest']
# Formula: Overlap ~ cohort * time * Hemisphere * L1L2 + print_base + speech_base +      (1 + L1L2 | subject)
# Data: IFG_all
# 
# REML criterion at convergence: -1432.1
# 
# Scaled residuals: 
#   Min      1Q  Median      3Q     Max 
# -2.9265 -0.6403 -0.1603  0.5629  5.5429 
# 
# Random effects:
#   Groups   Name        Variance  Std.Dev. Corr
# subject  (Intercept) 0.0103879 0.10192      
# L1L2L2      0.0005821 0.02413  0.10
# Residual             0.0248639 0.15768      
# Number of obs: 2184, groups:  subject, 101
# 
# Fixed effects:
#   Estimate Std. Error         df t value Pr(>|t|)    
# (Intercept)                                 1.966e-01  1.080e-02  9.948e+01  18.202  < 2e-16 ***
# cohortHeb L1                               -5.012e-02  1.086e-02  1.013e+02  -4.613 1.16e-05 ***
# timet3                                     -2.081e-02  3.604e-03  2.047e+03  -5.773 8.98e-09 ***
# HemisphereRight                            -5.258e-02  3.404e-03  1.970e+03 -15.447  < 2e-16 ***
# L1L2L2                                      2.019e-02  4.286e-03  1.005e+02   4.711 7.93e-06 ***
# print_base                                  1.285e-04  2.882e-05  1.009e+03   4.461 9.08e-06 ***
# speech_base                                 1.307e-04  2.933e-05  1.226e+03   4.457 9.07e-06 ***
# cohortHeb L1:timet3                        -2.783e-03  3.575e-03  2.043e+03  -0.778  0.43644    
# cohortHeb L1:HemisphereRight                1.401e-02  3.404e-03  1.970e+03   4.117 4.00e-05 ***
# timet3:HemisphereRight                      4.933e-04  3.404e-03  1.970e+03   0.145  0.88478    
# cohortHeb L1:L1L2L2                        -2.562e-02  4.205e-03  9.466e+01  -6.092 2.38e-08 ***
# timet3:L1L2L2                               7.774e-03  3.444e-03  2.068e+03   2.257  0.02411 *  
# HemisphereRight:L1L2L2                     -1.012e-02  3.404e-03  1.970e+03  -2.974  0.00298 ** 
# cohortHeb L1:timet3:HemisphereRight        -2.964e-04  3.404e-03  1.970e+03  -0.087  0.93063    
# cohortHeb L1:timet3:L1L2L2                  7.030e-03  3.445e-03  2.068e+03   2.041  0.04139 *  
# cohortHeb L1:HemisphereRight:L1L2L2         9.242e-03  3.404e-03  1.970e+03   2.715  0.00668 ** 
# timet3:HemisphereRight:L1L2L2              -3.259e-03  3.404e-03  1.970e+03  -0.958  0.33841    
# cohortHeb L1:timet3:HemisphereRight:L1L2L2 -2.728e-03  3.404e-03  1.970e+03  -0.802  0.42290    


# STG analysis ----
STG_ps_all <- readRDS("STG.Rda")
STG_lme <- lmer(data=STG_ps_all, STG ~ cohort*time*Hemisphere*L1L2+print_base+speech_base + (1+time+L1L2|subject))
summary(STG_lme)

#Linear mixed model fit by REML. t-tests use Satterthwaite's method ['lmerModLmerTest']
# Formula: STG ~ cohort * time * Hemisphere * L1L2 + print_base + speech_base +      (1 + time + L1L2 | subject)
# Data: STG_ps_all
# 
# REML criterion at convergence: -1749.6
# 
# Scaled residuals: 
#   Min      1Q  Median      3Q     Max 
# -3.9171 -0.4388 -0.0783  0.3057  5.7291 
# 
# Random effects:
# Groups   Name        Variance  Std.Dev. Corr       
# subject  (Intercept) 0.0032952 0.05740             
#          timet3      0.0018480 0.04299  -0.13      
#          L1L2L2      0.0003837 0.01959  -0.16 -0.24
# Residual             0.0020147 0.04489             
# Number of obs: 728, groups:  subject, 101
# 
# Fixed effects:
#                                              Estimate Std. Error         df t value Pr(>|t|)    
# (Intercept)                                 5.394e-02  6.177e-03  1.006e+02   8.733 5.64e-14 ***
# cohortHeb L1                               -9.689e-03  6.225e-03  1.028e+02  -1.556 0.122689    
# timet3                                     -7.689e-03  4.890e-03  9.560e+01  -1.572 0.119191    
# HemisphereRight                            -6.372e-03  1.678e-03  4.344e+02  -3.797 0.000167 ***
# L1L2L2                                     -8.750e-04  2.672e-03  1.051e+02  -0.327 0.743954    
# print_base                                 -8.082e-05  1.978e-05  5.192e+02  -4.085 5.11e-05 ***
# speech_base                                 8.303e-05  2.098e-05  5.780e+02   3.957 8.54e-05 ***
# cohortHeb L1:timet3                         1.449e-03  4.867e-03  9.462e+01   0.298 0.766612    
# cohortHeb L1:HemisphereRight                2.358e-03  1.678e-03  4.344e+02   1.405 0.160649    
# timet3:HemisphereRight                     -2.217e-03  1.678e-03  4.344e+02  -1.321 0.187247    
# cohortHeb L1:L1L2L2                        -9.611e-03  2.606e-03  9.748e+01  -3.688 0.000372 ***
# timet3:L1L2L2                              -5.029e-03  1.718e-03  4.709e+02  -2.928 0.003582 ** 
# HemisphereRight:L1L2L2                     -1.878e-03  1.678e-03  4.344e+02  -1.119 0.263743    
# cohortHeb L1:timet3:HemisphereRight         1.726e-05  1.678e-03  4.344e+02   0.010 0.991800    
# cohortHeb L1:timet3:L1L2L2                 -5.348e-04  1.718e-03  4.713e+02  -0.311 0.755776    
# cohortHeb L1:HemisphereRight:L1L2L2         2.925e-03  1.678e-03  4.344e+02   1.743 0.082100 .  
# timet3:HemisphereRight:L1L2L2              -2.000e-04  1.678e-03  4.344e+02  -0.119 0.905218    
# cohortHeb L1:timet3:HemisphereRight:L1L2L2  2.715e-03  1.678e-03  4.344e+02   1.618 0.106412

# IPC  analysis ----
IPC_ps_all <- readRDS("IPC.Rda")
IPC_lme <- lmer(data=IPC_ps_all, IPC ~ cohort*time*Hemisphere*L1L2+print_base+speech_base + (1+time+Hemisphere+L1L2|subject))
summary(IPC_lme)

# Linear mixed model fit by REML. t-tests use Satterthwaite's method ['lmerModLmerTest']
# Formula: IPC ~ cohort * time * Hemisphere * L1L2 + print_base + speech_base +      (1 + time + Hemisphere + L1L2 | subject)
# Data: IPC_ps_all
# 
# REML criterion at convergence: -769.2
# 
# Scaled residuals: 
#   Min       1Q   Median       3Q      Max 
# -2.08418 -0.54591 -0.03615  0.45302  2.63256 
# 
# Random effects:
# Groups   Name            Variance  Std.Dev. Corr             
# subject  (Intercept)     0.0148297 0.12178                   
#          timet3          0.0036802 0.06066  -0.23            
#          HemisphereRight 0.0021235 0.04608  -0.32  0.16      
#          L1L2L2          0.0008694 0.02949  -0.27 -0.14 -0.22
# Residual                 0.0075376 0.08682                   
# Number of obs: 728, groups:  subject, 101
# 
# Fixed effects:
#                                              Estimate Std. Error         df t value Pr(>|t|)    
# (Intercept)                                 1.853e-01  1.283e-02  9.735e+01  14.435  < 2e-16 ***
# cohortHeb L1                               -3.420e-02  1.291e-02  9.915e+01  -2.649  0.00941 ** 
# timet3                                     -1.671e-02  7.335e-03  8.450e+01  -2.278  0.02527 *  
# HemisphereRight                            -6.587e-02  5.689e-03  9.842e+01 -11.579  < 2e-16 ***
# L1L2L2                                     -1.391e-02  4.551e-03  1.058e+02  -3.057  0.00283 ** 
# print_base                                  3.258e-05  3.544e-05  4.443e+02   0.920  0.35832    
# speech_base                                 6.699e-05  3.749e-05  4.911e+02   1.787  0.07457 .  
# cohortHeb L1:timet3                        -9.735e-03  7.293e-03  8.366e+01  -1.335  0.18556    
# cohortHeb L1:HemisphereRight                9.723e-03  5.689e-03  9.842e+01   1.709  0.09056 .  
# timet3:HemisphereRight                      3.084e-03  3.329e-03  3.620e+02   0.926  0.35481    
# cohortHeb L1:L1L2L2                        -4.530e-03  4.425e-03  9.757e+01  -1.024  0.30853    
# timet3:L1L2L2                              -4.336e-03  3.300e-03  3.684e+02  -1.314  0.18973    
# HemisphereRight:L1L2L2                     -3.952e-03  3.246e-03  3.370e+02  -1.218  0.22425    
# cohortHeb L1:timet3:HemisphereRight         8.836e-03  3.329e-03  3.620e+02   2.654  0.00830 ** 
# cohortHeb L1:timet3:L1L2L2                 -2.099e-03  3.300e-03  3.686e+02  -0.636  0.52528    
# cohortHeb L1:HemisphereRight:L1L2L2         2.230e-03  3.246e-03  3.370e+02   0.687  0.49257    
# timet3:HemisphereRight:L1L2L2              -2.353e-04  3.246e-03  3.370e+02  -0.072  0.94226    
# cohortHeb L1:timet3:HemisphereRight:L1L2L2  2.228e-03  3.246e-03  3.370e+02   0.686  0.49298


# FFG analysis -----
FFG_ps_all <- readRDS("FFG.Rda")
FFG_lme <- lmer(data=FFG_ps_all, FFG ~ cohort*time*Hemisphere*L1L2+print_base+speech_base + (1+time+L1L2|subject))
summary(FFG_lme)

# Linear mixed model fit by REML. t-tests use Satterthwaite's method ['lmerModLmerTest']
# Formula: FFG ~ cohort * time * Hemisphere * L1L2 + print_base + speech_base +      (1 + time + L1L2 | subject)
# Data: FFG_ps_all
# 
# REML criterion at convergence: -1200.8
# 
# Scaled residuals: 
#   Min      1Q  Median      3Q     Max 
# -2.5518 -0.5346 -0.0666  0.3245  4.0251 
# 
# Random effects:
#   Groups   Name        Variance  Std.Dev. Corr       
# subject  (Intercept) 0.0060126 0.07754             
#          timet3      0.0012747 0.03570  -0.17      
#          L1L2L2      0.0004707 0.02170  -0.03  0.66
# Residual             0.0055402 0.07443             
# Number of obs: 728, groups:  subject, 101
# 
# Fixed effects:
#                                              Estimate Std. Error         df t value Pr(>|t|)    
# (Intercept)                                 8.728e-02  8.376e-03  9.595e+01  10.421   <2e-16 ***
# cohortHeb L1                               -1.830e-02  8.444e-03  9.809e+01  -2.167   0.0327 *  
# timet3                                     -1.251e-02  4.805e-03  9.036e+01  -2.603   0.0108 *  
# HemisphereRight                            -2.680e-02  2.783e-03  4.313e+02  -9.630   <2e-16 ***
# L1L2L2                                      5.126e-03  3.646e-03  1.002e+02   1.406   0.1628    
# print_base                                  4.252e-05  2.689e-05  4.203e+02   1.581   0.1146    
# speech_base                                 4.294e-06  2.784e-05  3.917e+02   0.154   0.8775    
# cohortHeb L1:timet3                        -6.899e-03  4.775e-03  8.950e+01  -1.445   0.1520    
# cohortHeb L1:HemisphereRight               -2.499e-03  2.783e-03  4.313e+02  -0.898   0.3697    
# timet3:HemisphereRight                      5.165e-03  2.783e-03  4.313e+02   1.856   0.0642 .  
# cohortHeb L1:L1L2L2                        -4.292e-03  3.558e-03  9.307e+01  -1.206   0.2308    
# timet3:L1L2L2                               2.394e-03  2.820e-03  4.753e+02   0.849   0.3963    
# HemisphereRight:L1L2L2                     -5.562e-03  2.783e-03  4.313e+02  -1.999   0.0463 *  
# cohortHeb L1:timet3:HemisphereRight        -1.744e-04  2.783e-03  4.313e+02  -0.063   0.9501    
# cohortHeb L1:timet3:L1L2L2                 -3.051e-03  2.820e-03  4.755e+02  -1.082   0.2799    
# cohortHeb L1:HemisphereRight:L1L2L2         3.897e-03  2.783e-03  4.313e+02   1.400   0.1621    
# timet3:HemisphereRight:L1L2L2               3.714e-05  2.783e-03  4.313e+02   0.013   0.9894    
# cohortHeb L1:timet3:HemisphereRight:L1L2L2  1.296e-03  2.783e-03  4.313e+02   0.466   0.6417
```
